# Influenza A virus reassortment is strain dependent

**DOI:** 10.1101/2022.09.26.509616

**Authors:** Kishana Y. Taylor, Ilechukwu Agu, Ivy José, Sari Mäntynen, A.J. Campbell, Courtney Mattson, Tsui-wen Chou, Bin Zhou, David Gresham, Elodie Ghedin, Samuel L. Díaz Muñoz

## Abstract

RNA viruses can exchange genetic material during coinfection, an interaction that creates novel strains with implications for viral evolution and public health. Influenza A viral genetic exchange occurs when genome segments from distinct strains reassort in coinfection. Predicting potential reassortment between influenza strains is a longstanding goal. Experimental coinfection studies have shed light on factors that limit or promote reassortment. However, determining the reassortment potential between diverse Influenza A strains has remained an elusive goal. To fill this gap, we developed a high throughput genotyping approach to quantify reassortment among a diverse panel of human influenza virus strains, encompassing 41 years of epidemics, multiple geographic locations, and both circulating human subtypes A/H1N1 and A/H3N2. We found that the reassortment rate (proportion of reassortants) is an emergent property of a pair of strains where strain identity is a predictor of the reassortment rate. We show little evidence that antigenic subtype drives reassortment as intersubtype (H1N1xH3N2) and intrasubtype reassortment rates were, on average, similar. Instead, our data suggest that certain strains bias the reassortment rate up or down, independently of the coinfecting partner. We also observe that viral productivity is an emergent property of coinfections and that it is not correlated to reassortment rate, thus affecting the total number of reassortant progeny produced. Assortment of individual segments among progeny, and pairwise segment combinations within progeny, were not random and generally favored homologous combinations. This outcome was not related to strain similarity or shared subtype. Reassortment rate was closely correlated to both the proportion of unique genotypes and the proportion of progeny with heterologous pairwise segment combinations. We provide experimental evidence that viral genetic exchange is potentially an individual social trait subject to natural selection, which implies the propensity for reassortment is not evenly shared among strains. This study highlights the need for research incorporating diverse strains to discover the traits that shift the reassortment potential as we work towards the goal of predicting influenza virus evolution resulting from segment exchange.

## Introduction

RNA viruses can interact during coinfection to exchange genetic material, creating novel strains that can evade immunity, and impact host range. Coinfection of influenza viruses provides the opportunity for their negative-sense RNA genome segments to be swapped during virion assembly, a process called reassortment (reviewed in: Lakdawala et al., 2016; McDonald et al., 2016; Steel and Lowen, 2014). Influenza A pandemics have been sparked by the emergence of reassortant strains (Medina and García-Sastre, 2011), most recently with the 2009 H1N1 pandemic strain (Garten et al., 2009). In the case of human seasonal influenza, every season there is a large collection of co-circulating lineages (Ghedin et al., 2005; Holmes et al., 2005; Nelson et al., 2008a; Rambaut et al., 2008) that can reassort (Holmes et al., 2005; Nelson et al., 2008b, 2008a; Rambaut et al., 2008), creating novel variants that have, for instance, spread antiviral resistance to adamantane before its widespread use (Simonsen et al., 2007). Thus, reassortment has remained a topic of interest due to its substantial public health implications and consequences on the global epidemiology of influenza A viruses.

Predicting the outcomes of coinfection between intact strains (i.e. not experimentally manipulated, wild type) and potential reassortment is a longstanding goal of basic research and public health risk assessment. Experimental coinfection studies provide a powerful tool to assess coinfection outcomes and dissect patterns of segment exchange (Steel and Lowen, 2014) and have made substantial progress in identifying factors that limit or promote reassortment. The first experimental coinfection study (Lubeck et al., 1979) showed non-random association between segments in a A/PR/8/1934(H1N1) and A/Hong Kong/1/68(H3N2) coinfection, particularly in lower-than-expected exchange among the three segments that form the polymerase complex. Subsequent studies employing variations on the experimental coinfection approach supported non-random associations of segments between divergent heterosubtypic strains (Essere et al., 2013; Greenbaum et al., 2012; Li et al., 2008). These results stimulated research into the factors that may restrict segment exchange (White and Lowen, 2018) including: the replicative capacity of coinfecting viruses (Octaviani et al., 2010); functional incompatibilities between proteins (Li et al., 2008); and RNA-RNA interactions involving genome packaging signals (Essere et al., 2013; White et al., 2019; White and Lowen, 2018).

In contrast, several researchers argued that reassortment among intact strains was more pervasive than implied by these initial experimental coinfection studies (Brooke, 2017; Koelle et al., 2019; Marshall et al., 2013; Steel and Lowen, 2014). Extending the conclusions of these studies to determining the reassortment potential of intact strains was hampered by methodological details, such as: using reverse genetics to “force” genotype combinations that may not occur with intact viruses; allowing multi-cycle replication, which confounds the effects of between-cell competition on fitness with the intracellular process of reassortment; and examining a small sample of progeny viruses. These limitations were overcome by a landmark study conducted by Marshall et al. (Marshall et al., 2013) which showed that near-identical strains (differing only in synonymous substitutions to facilitate genotyping) of A/Panama/2007/99(H3N2) exhibited reassortment at levels approaching random expectations (Marshall et al., 2013). A second major contribution was tight control of infection conditions that limited infections to a single cycle, thus isolating reassortment outcomes from between-cell competition. This system was then leveraged to study reassortment of A/Panama/2007/99(H3N2) and A/Netherlands/602/2009(H1N1pdm), while controlling for homologous (self) reassortment. This study revealed that reassortment was efficient between strains of different subtypes, but that segment exchange was non-random; bias towards specific segments or segment combinations (e.g. homologous PB2-PA combinations) were detected in the progeny. Thus, even though the *rate* of reassortment may not differ between strains, the specific combinations of segments in the progeny can differ substantially. This result underscored that determining reassortment potential cannot be limited to a single measure. A number of studies have incorporated an increasingly detailed accounting of reassortment potential (Ganti et al., 2021; Phipps et al., 2017) including: i) the proportion of progeny that are reassortant; ii) the number of genotypes produced; ii) the relative fitness of segments; and iii) the pairwise linkage of segments and other population genetic measures such as evenness and entropy. In sum, two conclusions regarding human influenza A have become widely accepted in the literature: First, that very similar strains generate a high proportion of reassortant progeny and a closer-to-random segment exchange; and, second, that divergence between parental viruses generates biased segment combinations.

Despite much progress, determining the reassortment potential between diverse influenza A strains has remained an elusive goal. Due to methodological limitations, studies to date have largely tested one strain pair at a time while studies involving more than two viral backgrounds are rare. Limited experimental studies have been done between diverse human seasonal strains spanning decades. This is crucial for three reasons. First, the factors that limit or favor reassortment can be biased by the selection of strains used to determine reassortment potential. Second, reassortment potential between strains may change as the virus evolves. Third, from an evolutionary perspective, while two viruses are required for genetic exchange, each may differ in their propensity to exchange genes (Díaz-Muñoz et al., 2013; O’Keefe et al., 2010), as observed in individual variability in meiotic recombination rate between animals (Coop & Przeworski, 2007, Ritz et al. 2017, Smukowski & Noor 2011). More generally, in nature there is ample potential for diverse intact strains to encounter each other and reassort, underscoring the need to incorporate this variability into our experimental design (Díaz-Muñoz, 2019).

To fill this gap, we developed a high throughput genotyping approach to facilitate quantification of influenza A virus reassortment between human strains. This tool allowed us to examine coinfection outcomes among a diverse collection of strains, representing multiple epidemics; geographic origins; and both circulating human influenza A subtypes, H3N2 and H1N1. Our overall objectives were to test some of the conclusions regarding reassortment potential in the literature and address questions that remain intractable with current experimental approaches. Specifically, we aimed to: 1) use a statistically robust sample size to test strain traits (genetic similarity, subtype, epidemic season) that may influence reassortment potential; 2) provide a detailed accounting of coinfection outcomes (replication, proportion of reassortants, relative fitness of segments, segment linkage patterns, and genotype diversity); and 3) examine associations between coinfection outcomes (e.g. testing if viral replication of the coinfection is correlated with the reassortment rate).

We found a wide range of reassortment outcomes emerging from the pairwise strain combinations involved in the experimental coinfections. Of note, we find that high reassortment rates and random patterns of segment exchange were possible between divergent or heterosubtypic strains. Furthermore, there was no significant relationship between genetic similarity and various measures of reassortment. We show that viral kinetics vary independently of the reassortment rate, affecting the total number of reassortants produced by each coinfection. Most significantly, we found evidence of strain-dependent variation in reassortment wherein some strains bias reassortment rate upwards, regardless of the coinfecting partner. This result suggests that reassortment potential may be strain specific and may be a trait of individual virus strains.

## Methods

### Cells and Viruses

We maintained MDCK (Madin-Darby Canine Kidney) cells in minimum essential media supplemented with 5% FBS (Fetal Bovine Serum). Cells were checked for mycoplasma contamination by PCR. We obtained five wild type influenza A virus strains, a generous gift from the lab of Dr. Ted Ross, which we abbreviate here as: CA09, HK68, PAN99, SI86, TX12 (Table 1). These stocks were originally egg passaged and we made stocks through low multiplicity of infection (MOI) propagation in MDCK cells (obtained from ATCC/BEI), thus these virus stocks are a diverse population with mutations and internal deletions as is characteristic of natural (Saira et al., 2013) and lab (Xue et al., 2016) influenza viral stocks, even to some extent those created via reverse genetics (Russell et al., 2019).

**Table 1.** Influenza A strains used in experiments and their properties.

| Strain Name | Subtype | Year | Isolation | Abbreviation |
| --- | --- | --- | --- | --- |
| A/Hong Kong/1/1968(H3N2) | H3N2 | 1968 | Hong Kong | HK68 |
| A/Singapore/1986(H1N1) | H1N1 | 1986 | Singapore | SI86 |
| A/Panama/2007/1999(H3N2) | H3N2 | 1999 | Panama | PAN99 |
| A/California/07/2009(H1N1)pdm | H1N1 | 2009 | California | CA09 |
| A/Texas/2012(H3N2) | H3N2 | 2012 | Texas | TX12 |

### Infections and Plaque Assays

#### Pairwise Infections

We conducted infections following the conditions of Marshall et al. (2013). Briefly, a 6-well plate with 1 X 10^6^ of MDCK cells was coinfected with a pair of strains in equal proportion at a MOI of 10. We synchronized infections at 4°C for 1 hour to promote equal attachment of viruses. To limit infections beyond one cycle of replication, we conducted infections in the absence of trypsin (Klenk et al., 1975; Tobita et al., 1975) collected viral supernatants at 12hr post infection and stored them at −80°C for subsequent genotyping and titration.

#### Control Infections

We conducted several control experiments to test the genotype by barcode sequencing approach (see below). First, we made biological replicates of the same coinfections for three strain pairs (HK68xCA09 n = 2, and HK68xPAN99 n = 2, and CA09/PAN99 n=3). Second, we conducted infections of HK68xCA09 at increasing MOI’s (0.01, 1, and 10), which are expected to lead to increasing levels of coinfection and reassortment (Marshall et al. 2013). Finally, we evaluated the parental stocks used to initiate coinfections to evaluate the genotyping pipeline and establish thresholds for assigning sequencing reads to specific strains.

To isolate clonal populations for genotyping progeny from the infections, we conducted plaque assays. Plaques were picked with 1mL sterile pipette tips and placed in DPBS in deep-well 96-position plates for genotyping (see below). We also conducted plaque assays from the same supernatants to quantify infectious particle production yield of experimental coinfection supernatants.

### Genotype by Barcode Sequencing

To ascertain the eight-segment genotype of the viral progeny (i.e. isolated plaques representing clonal populations), we used a strategy we term **G**enotype **b**y **B**arcode **Seq**uencing (GbBSeq), inspired by Bar-seq experiments of yeast mutant libraries (Robinson et al. 2014). We describe the strategy in detail in Supplementary Material. Briefly, the picked plaques are used as RNA template in a one-step RT-PCR reaction with: 1) a uni13 primer (targeting a conserved region shared by all influenza A virus segment ends) barcoded to correspond with each of 96 wells; and 2) 10 primers (6 for internal segments and 4 primers customized for H1/H3 and N1/N2 subtypes), targeting the internal regions of genome segments barcoded to correspond with each coinfection (represented by a plate). These barcoded PCR products were pooled, diluted, and used as template in a second PCR reaction that added adapters for paired-end sequencing on the Illumina MiSeq platform. All GbBSeq experiments in this paper (including controls) were conducted in a single MiSeq run. This protocol bears some similarity to the genotyping-in-thousands approach or GT-seq (Campbell et al., 2015), but was independently conceived by our research group prior to the cited publication.

#### Data analysis pipeline

The analysis pipeline is posted on GitHub (https://github.com/sociovirology/human_influenza_GbBSeq). In brief, paired-end reads were demultiplexed and trimmed using CUTADAPT (v2.6), allowing only exact matches, resulting in files that contained reads for a single clonal population from a given coinfection. We conducted no further quality filtering, as quality scores were very high after read trimming and read merging using PEAR (v0.9.6). We matched the reads of each file to a database containing the full-length amplicons for strains included in each cross, using the usearch_global command at 98% percent nucleotide identity in USEARCH (v8.1.1861). The output of this database search was a file for each clonal progeny, which includes the match, PID, mismatches, and length of alignment, as well as other features, output for analysis in R (v3.4.1).

#### Genotype Assignment

To assign the full-length genotype of a progeny, each of the eight segments was assigned to one or another parental strain. Because progeny plaques were clonal populations and each segment was represented by multiple reads, we tallied the number of database matches and used a majority rule (i.e. consensus) to assign segments to a strain.

### Reassortment quantification and statistics

We scored individual progeny as reassortant if they contained one or more segments from different parental strains. We calculated the reassortment <u>rate</u> between two strains in a coinfection by calculating the percentage of reassortant progeny resulting from a given coinfection (proportion reassortant = reassortant plaques / total plaques isolated). We calculated deviations from random expectations in the reassortment rate using a Fisher exact test (eight segments in 256 unique combinations, 2 of which are parents; 254/256 = 99.22%). Because coinfections vary in the number of infectious progeny they generate (i.e. some coinfections are more productive in terms of viral replication), we estimated the *total* number of reassortants by determining the final titer of the supernatants (the same supernatant used to isolate plaques for genotyping) and multiplying by the respective reassortment rate. We then set out to determine whether individual strains could influence the reassortment rate of a coinfection, analogous to the general combining ability (Henderson, 1952) on the recombination rate in animals (Coop and Przeworski, 2007; Ritz et al., 2017; Smukowski and Noor, 2011). While any given strain could have low or high reassortment rates, the strain could tend to generate higher or lower reassortment rates when controlling for the influence of the coinfecting partner. To determine whether particular strains had a tendency to increase or decrease reassortment, we used analysis of variance to calculate the individual contribution of each strain to the reassortment rate.

### Quantification of genotypes generated by experimental coinfections

To estimate the unique genotypes produced by each experimental coinfection, we determined the eight-segment composition of each plaque and calculated genotype richness (S): the number of unique eight-segment combinations. To control for differences in the number of total plaques with complete eight-segment genotypes (N’) isolated in each coinfection, we calculated the proportion of unique genotypes (S / N’).

### Segment bias calculations

To calculate bias in the genotype segment composition according to its parental origin, we examined the proportion of parental origin in each individual segment (across a given experimental coinfection) and used a Fisher exact test to examine deviations from the null frequency of 0.50.

### Pairwise interactions among segments

We conducted pairwise analyses of segment associations and calculated linkage disequilibrium statistics to determine whether segment exchange was free among parental segments or whether segments were linked with respect to their parental origin. There are 28 possible segment pair combinations among eight segments. For each of these segment pair combinations, there are four potential combinations of the two segments derived from two parental strains: two of which are homologous and two heterologous. We calculated the proportion of plaque isolates that carried heterologous segment combinations for each for the 28 possible segment combinations, in each experimental coinfection.

### Effect of sampling on estimates of reassortment and segment bias

To determine how close experimental coinfections came to the 256 theoretically possible segment combinations, we conducted simulations to test the proportion of all possible segment combinations with the sample of 94 reads. Additionally, we used repeated experimental coinfections of select strain combinations to conduct simulations using empirical data to test the effect of sampling on the number of recovered genotypes.

## Results

### Genotype by Barcode Sequencing: a system for high-throughput quantification of reassortment between diverse strains

We used GbBSeq to ascertain the eight-segment genotypes of viral progeny derived from multiple experimental coinfections between our diverse panel of human influenza A strains. We sequenced 16 experimental coinfections (including control and replicate infections) in one MiSeq sequencing run resulting in a total of 25,729,649 reads, of which 15,501,869 were demultiplexed and yielded matches to the strain database. These reads covered 1,536 clonal progeny (mean reads / cross = 968,866 reads, range 219,326 – 1,544,248). The average read depth for each segment was 1,293x and was lowest for two polymerase complex segments (PB2 = 45.6x, PB1 = 16.3x), which are subject to large internal deletions, a well-known phenomenon (Russell et al., 2019; Xue et al., 2016). These differences in coverage resulted in incomplete genotypes, primarily involving segment PB1. We established a majority rule for strain assignment (e.g. >50%), but on average strains were assigned at a higher proportion, 0.945 ± 0.110; overall, 90% of segments were assigned to a strain by over 75% of reads. Reassortment rate estimates (i.e. proportion of genotyped progeny with segments from >1 parent) were repeatable among biological replicates in experimental coinfections, regardless of whether the coinfection generated low, medium, or high reassortment rates (**Figure 1A**). On average, replicate reassortment rate estimates differed by ±0.06246 (range = 0.0100 – 0.1143). Reassortment rate increased as a function of MOI and the data fit within the standard error of a model of exponential increase in reassortment as the MOI increased (**Figure 1B**), as previously reported (Marshall et al., 2013).

**Figure 1.**
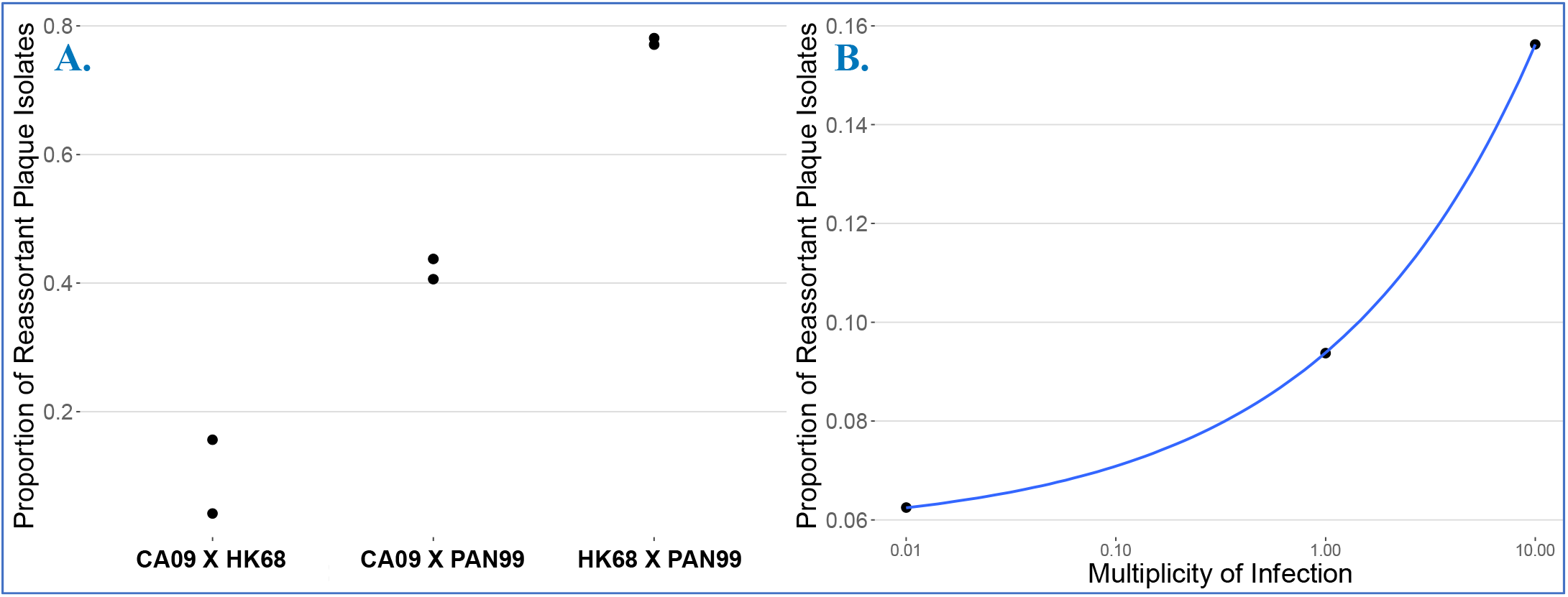
Genotype by Barcode Sequencing is replicable and measures reassortment accurately. **A**. The proportion of reassortant plaque isolates in biological replicates of experimental coinfections initiated at MOI 10. The insets show 8 segment genotype composition of isolated plaques in these replicates representing coinfections with low, medium, and high rates of reassortment. **B**. The proportion of reassortant plaque isolates from experimental coinfections at three multiplicities (0.01, 1, 10) of infection for the CA09 x HK68 experimental coinfection. The fray shading around the blue line represents the standard error around an exponential distribution.

### Reassortment is an emergent property of a pair of strains

We set out to investigate reassortment potential among five human influenza A strains (CA09, HK68, PAN99, SI86, TX12 see **Table 1**) encompassing 41 years of epidemics, multiple geographic locations, and both circulating human subtypes. We conducted all possible pairwise experimental coinfections between these strains (n = 10) in MDCK cells at an MOI of 10 and determined the genotypes of progeny isolated via plaque assay. Genotypes for individual progeny are depicted as rows in each panel (**Figure 2A**), with columns indicating the genotype (i.e. parental origin) for each of the eight segments.

**Figure 2.**
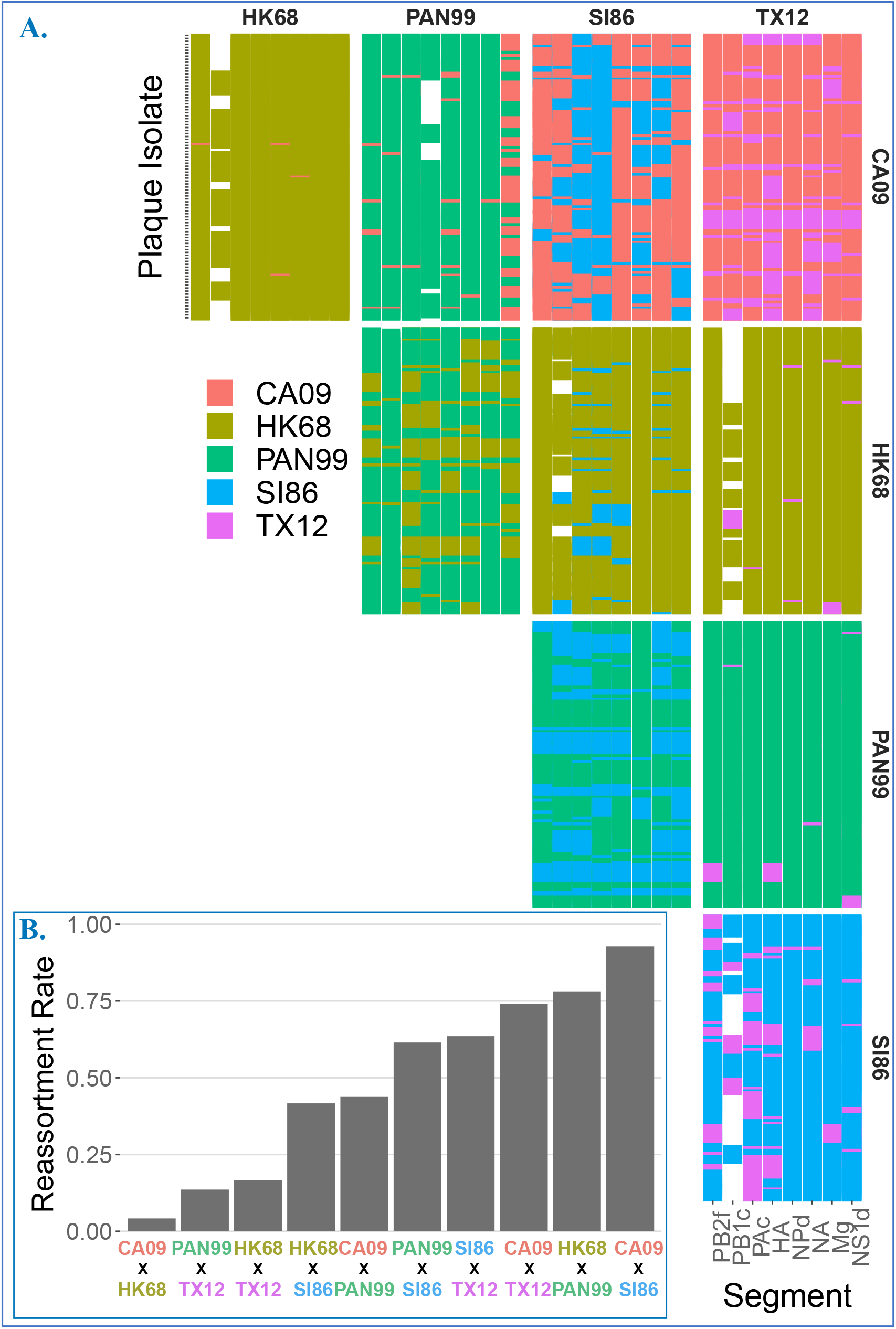
Reassortment plots and rates determined from plaque isolates. **A**. Reassortment plot of genotyped progeny isolated from each coinfection performed in this study. Genotypes for individual progeny are depicted as rows in each panel with columns indicating the genotype (i.e. parental origin) for each of the eight segments. White spaces show genotypes where one or more segments were not identified or recovered. **B**. Reassortment rates (proportion of reassortant plaque isolates) for each coinfection. Red line indicates theoretical maximum reassortment, gray line indicates 40% reassortant progeny

From this primary data, we initially measured and analyzed the reassortment rate, defined as the proportion of progeny harboring at least one segment from a different parent. All pairwise experimental coinfections (n = 10) had measurable reassortment rates (**Figure 2A**). Reassortment rate estimates did not differ substantially, qualitatively or quantitatively, when plaque isolates with incomplete genotypes were included or excluded. Thus, reassortment rates are presented using all data, which is a conservative approach because inclusion of genotypes that are missing segments decrease the chance of detecting a reassortant. Reassortment rate ranged from 92.71% to 4.17% with an average of 48.95% ± 30.08 (mean ± sd). Overall, 7/10 coinfections generated reassortment rates of >40% (**Figure 2B**). Under the assumption of completely free reassortment, the percentage of reassortants expected is 99.22%, i.e. 254/256. One experimental coinfection approached this expectation numerically, CA09xSI86 (92.71 %), but like all other coinfections, random expectations for reassortment rate were rejected (Exact binomial: p < 0.001).

To test if the reassortment rates calculated for each experimental coinfection (represented by gray bars in **Fig 2B**) differed statistically, we determined whether reassortment rates were equal. A test for equal proportions suggested that reassortment rates (e.g. values depicted in bars in **Figure 2B**) for experimental coinfections were not equal (p < 0.001). Specifically, a pairwise comparison of proportions revealed 29/45 of the pairwise rate comparisons were statistically significantly different (Holm adjusted p < 0.05). For instance, the reassortment rate between the SI86xTX12 (0.635) and PAN99xSI86 (0.615) coinfections were not statistically different, but both were different from CA09xSI86 (0.927). These differences were not explained by genetic similarity (Adj. R^2^ =−0.07118, p = 0.5438; **Table 2**, Supplementary **Figure S2** and **Table S1**) or subtype (**Figure 3**, below), suggesting a distinct reassortment rate emerged from each strain combination (**Figure 2A**).

**Table 2.**
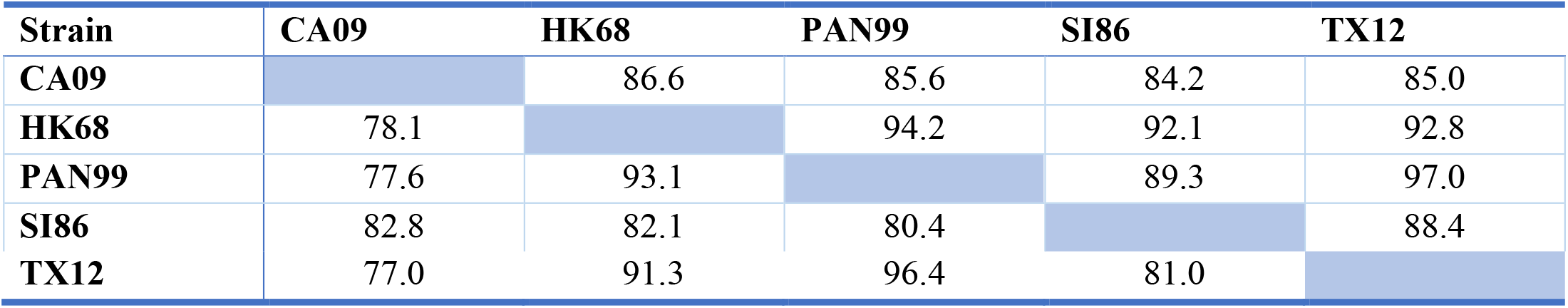
Pairwise percent nucleotide identity among Influenza A strains used in experimental coinfections. Values above the diagonal exclude antigenic segments HA and NA which are more highly variable than the internal segments.

**Figure 3.**
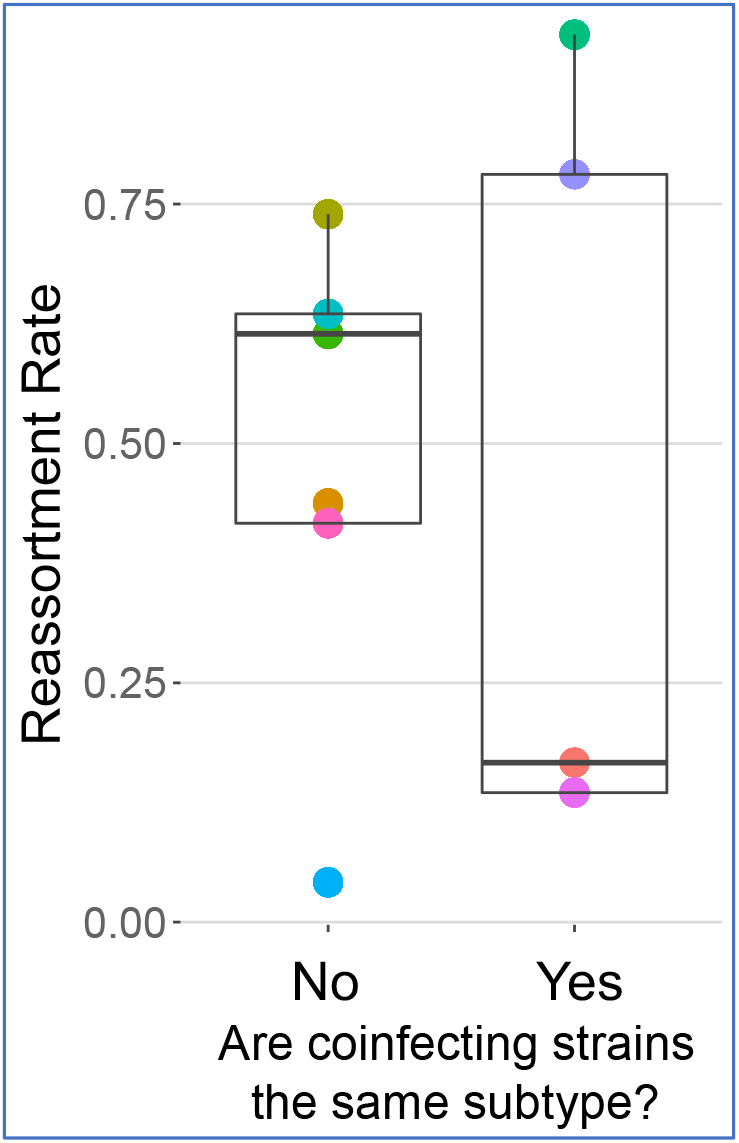
There is no significant difference in reassortment rates in intrasubtype (H1N1 or H3N2) coinfections versus intersubtype coinfections (H1N1xH3N2).

### Inter- and intrasubtype coinfections do not differ in reassortment rate

Experimental (Essere et al., 2013; Greenbaum et al., 2012; Li et al., 2008) and epidemiological (Nelson and Holmes, 2007) evidence has led to the idea that the subtype of coinfecting influenza A strains can influence reassortment, and specifically that intrasubtype reassortment should be more common (Nelson et al., 2008b, 2006). To determine whether antigenic subtype influenced reassortment rate, we examined reassortment rates within and between H3N2 and H1N1 subtypes. The mean intrasubtype reassortment rate was 0.50 ± 0.410 and the mean for coinfections between different subtypes (i.e. H3N2/H1N1) was 0.48 ± 0.248. There was no statistically significant difference in mean reassortment rates (t = 0.095, p = 0.9285) of inter- vs. intrasubtype coinfections (**Figure 3**).

### Reassortment rates are strain-dependent

Given that reassortment rate appeared to be an emergent property of each strain combination and did not relate to genetic similarity (Table 2) or subtype (Figure 3), we set out to determine whether individual strains could influence the reassortment rate of a coinfection and thus represent a trait of individual viruses, as observed for recombination rate in animals (Coop and Przeworski, 2007; Ritz et al., 2017; Smukowski and Noor, 2011). Strains generated a range of reassortment rates (Fig 2) and we were further interested in testing whether each strain consistently biased reassortment rate (upwards on downwards), independent of the coinfecting partner. We examined the average proportion of reassortants in each experimental coinfection (**Figure 4A**) using ANOVA removing the intercept from the regression, reflecting the biological reality that there is no reassortment if there is no coinfection partner. This analysis showed individual strains were a statistically significant predictor of reassortment rate (ANOVA: F = 15.75, df = 5, p = 0.004). Model coefficients (**Figure 4B**) represent a statistical calculation of the average individual contribution to the reassortment rate, in the theoretical absence of another strain. Specifically, the analysis suggested PAN99, HK68, and SI86 strains tended to, on average, increase reassortment, regardless of which other strain was coinfecting. For instance, SI86 had a significant ANOVA coefficient of 0.345; across the dataset the average reassortment rate for coinfections including SI86 was 0.648, with no coinfection below 0.416. In contrast, CA09 did not have a significant coefficient and exhibited the widest range of coinfection rates (0.417 – 0.927), indicating it did not influence the reassortment rate consistently. These results suggest that particular strains can significantly push the reassortment rate up or down. Thus, reassortment rates are an emergent property of the strain combination but can also be biased towards more or less reassortment by particular strains.

**Figure 4.**
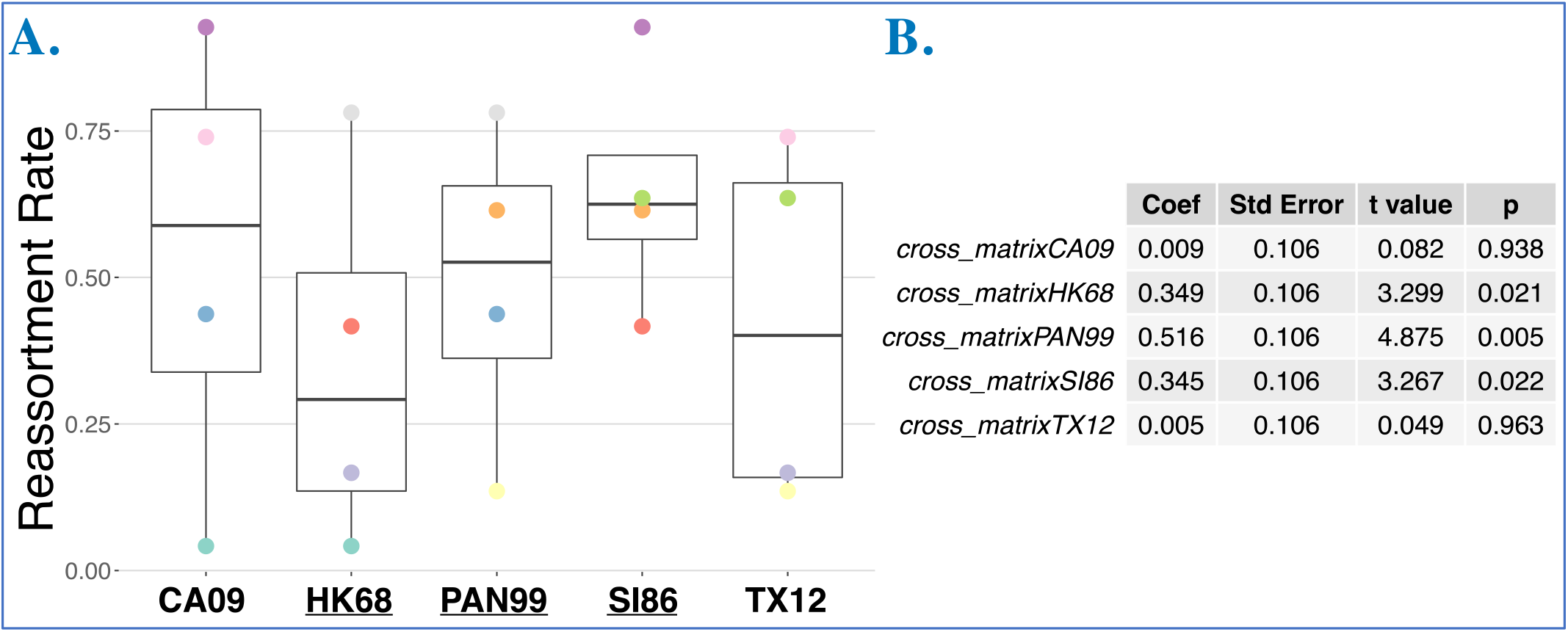
Reassortment rate by strain. **A**. Reassortment rates by strain, calculated from genotyping plaque isolates in ten experimental coinfections. Because reassortment rates are a pairwise property the same reassortment rate (points) appears in multiple strains, this is indicated by a shared color. **B**. Coefficients of an ANOVA model, showing that strain is a significant predictor of reassortment rate, with coefficients indicating individual strain contribution to reassortment rate.

### Viral yield affects the total number of reassortant progeny produced

Multiple studies suggest that cellular coinfection can change influenza A virus production in single-strain infections (Isken et al., 2012; Jacobs et al., 2019; Martin et al., 2020; Phipps et al., 2020), a phenomenon termed multiplicity dependence (Phipps et al., 2020). We reasoned that the mixed strain coinfections we conducted would affect viral reproduction of each coinfection. Thus, to gain insight into whether viral infection kinetics can affect the *total number* of reassortant progeny, we measured supernatant titers of each experimental coinfection. The titer of each coinfection (**Figure 5A**, *top bars*) was not correlated with the reassortment rate (**Figure 5B**). To estimate the total number of reassortant progeny, we multiplied the titer by the reassortant rate for each coinfection (**Figure 5A**, *bottom*) and show that the rank order of reassortment is different depending on whether total reassortants or reassortment rate is used.

**Figure 5.**
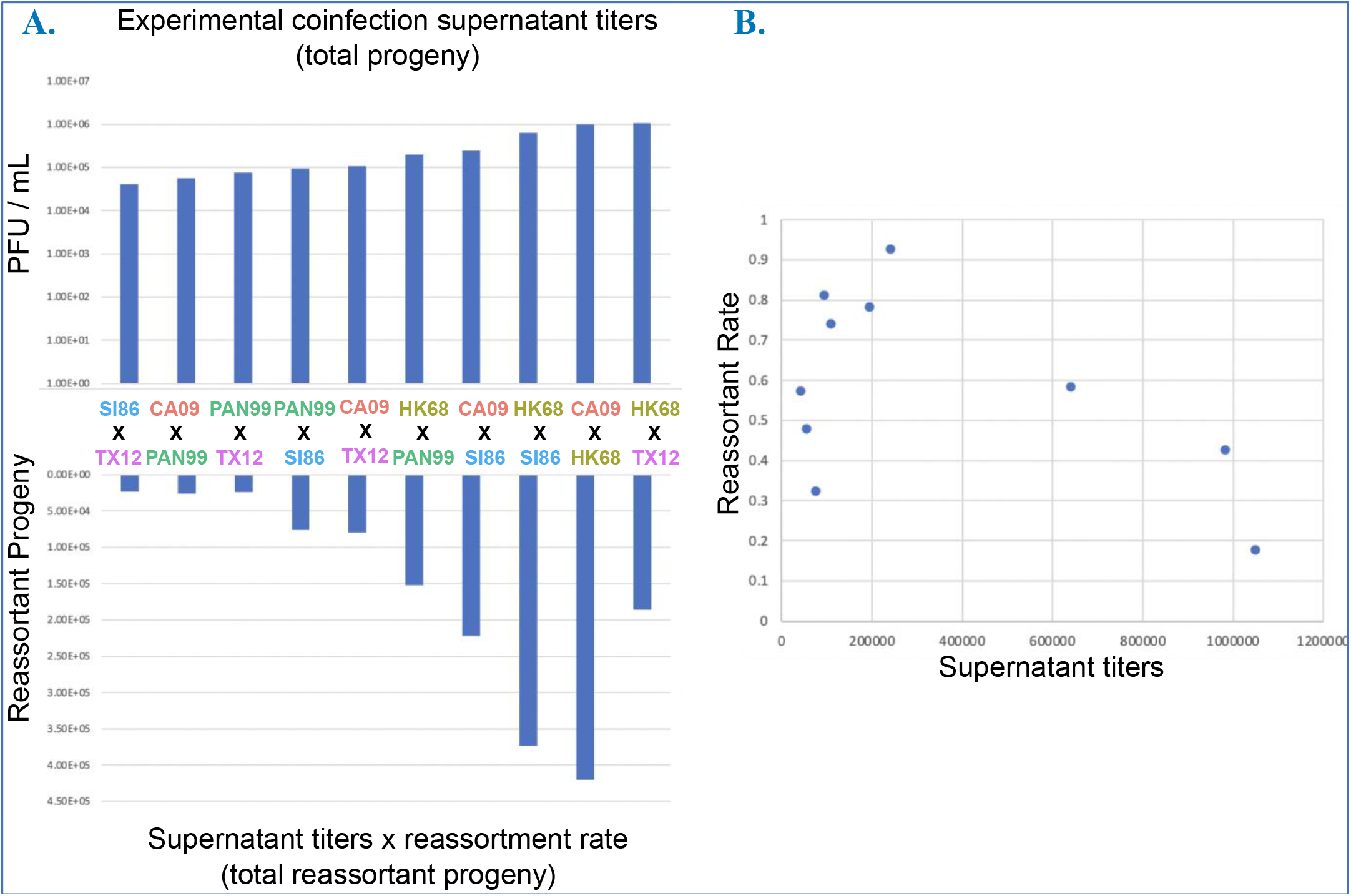
Viral kinetics of coinfections affect the total production of reasortants. **A**. Top bars show final infectious titers (PFU/mL) of each coinfection, in ascending order. Bottom bars show an estimate of the total reassortants produced by the coinfection, by multiplying the corresponding reassortment rate for each coinfection with its titer. Note that the rank order of total progeny (top bars), and total reassortant progeny (bottom bars), and reassortment rate (Figure 2B) is different. **B**. Reassortment rate is not correlated with titers of the same experimental coinfection.

### Many segments contribute to reassortment and individually deviate from random representation in progeny with respect to strain

Because reassortant progeny are defined as having one or more genome segments from different parental viruses, reassortment rates are not necessarily informative about patterns of segment exchange. To gain a segment-centric view on reassortment, we examined genotype representation across all progeny isolated from an experimental coinfection. Under unbiased coinfection conditions, for each of the eight IAV genome segments there is a 50/50 chance of either of the two parental segments being incorporated into a progeny virion, via random assortment. Biases in the parental strain origin of each segment, or deviations from random expectations, indicate potential differences in the relative fitness of segments during incorporation into progeny virions and coinfection. Upon examining the progeny across all coinfections, we first found that, on average, 7.100 ± 1.595 out of the eight segments were involved in reassortment, i.e. each strain contributed at least one of its segment alleles among all isolated plaques (**Figure 6A**). However, segment alleles were usually statistically biased towards one parental strain (**Figure 6B**, binomial exact test for deviation from 0.5 proportion at p < 0.05). Across all coinfections, only 13/80 segments conformed to random assortment (**Fig. 6B**; Supplementary Table S2). One intersubtype coinfection, SI86xPAN99, accounted for five of these freely assorting segments, with 6/8 segments adhering to random assortment in this cross. There was no relationship between the reassortment rate and the number of segments with parental strain representation bias (Supplementary Figure S2). Analyzing only reassortant plaques did not qualitatively change the results (Supplementary Figure S3).

**Figure 6.**
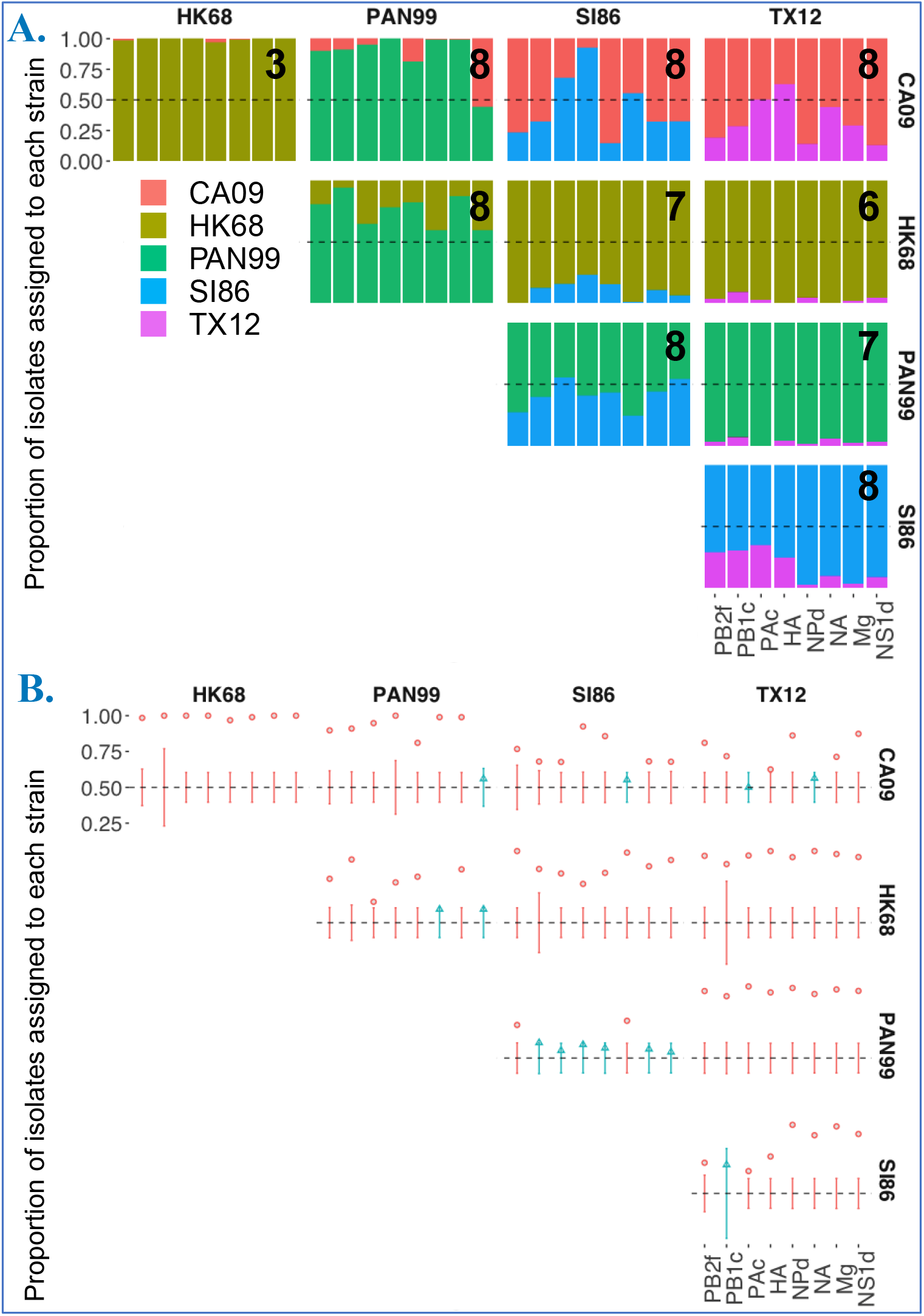
Many segments participate in genetic exchange and are non-randomly distributed in progeny plaque isolates with regard to strain. **A**. Plot shows frequency of each strain’s allele for each segment in plaque isolates. The number on the top right corner of each plot indicates the number of segments that participated in reassortment. **B**. Depicts which segment frequencies are within (blue points) or outside (red points) the confidence interval for 50:50 distribution of strain alleles among plaque isolates.

### Non-random pairwise segment associations were not limited to divergent strains and correlated with reassortment rate

Under completely free reassortment, segments randomly assort. Non-random pairwise associations between segments (i.e. linkage) have been observed, particularly for divergent strains. Using 2 influenza strains in a coinfection, there are 4 possibilities for pairwise segment combinations with each expected to be represented in 25% of progeny if there is no bias in pairwise segment associations (for example, in a coinfection between HK68 and SI86 the HA and NA segments could assort in the following ways: HA_HK68_ + NA_HK68_, HA_HK68_ + NA_SI86_, HA_SI86_ + NA_HK68_, HA_SI86_ + NA_SI86_). Two of these combinations have both segments derived from the same parent (i.e. homologous) and the remaining two have segments derived from each parent (i.e. heterologous). The latter is diversity generating, representing new combinations of segments. To simplify pairwise data and examine broad reassortment potential, we focused our analysis on quantifying the heterologous versus homologous plaques in each pairwise segment combination (out of 28 possible with 8 segments). Overall, the average proportion of plaques with heterologous pairwise combinations was significantly different according to the strain combination in each experimental coinfection (**Figure 7A**, ANOVA F =14.698, p < 0.0001). This measure did not vary according to the strain similarity or subtype of each coinfection. The reassortment rate was correlated to the average proportion of heterologous plaques (Adj. R^2^: 0.505, p =0.0128; **Figure 7B**).

**Figure 7.**
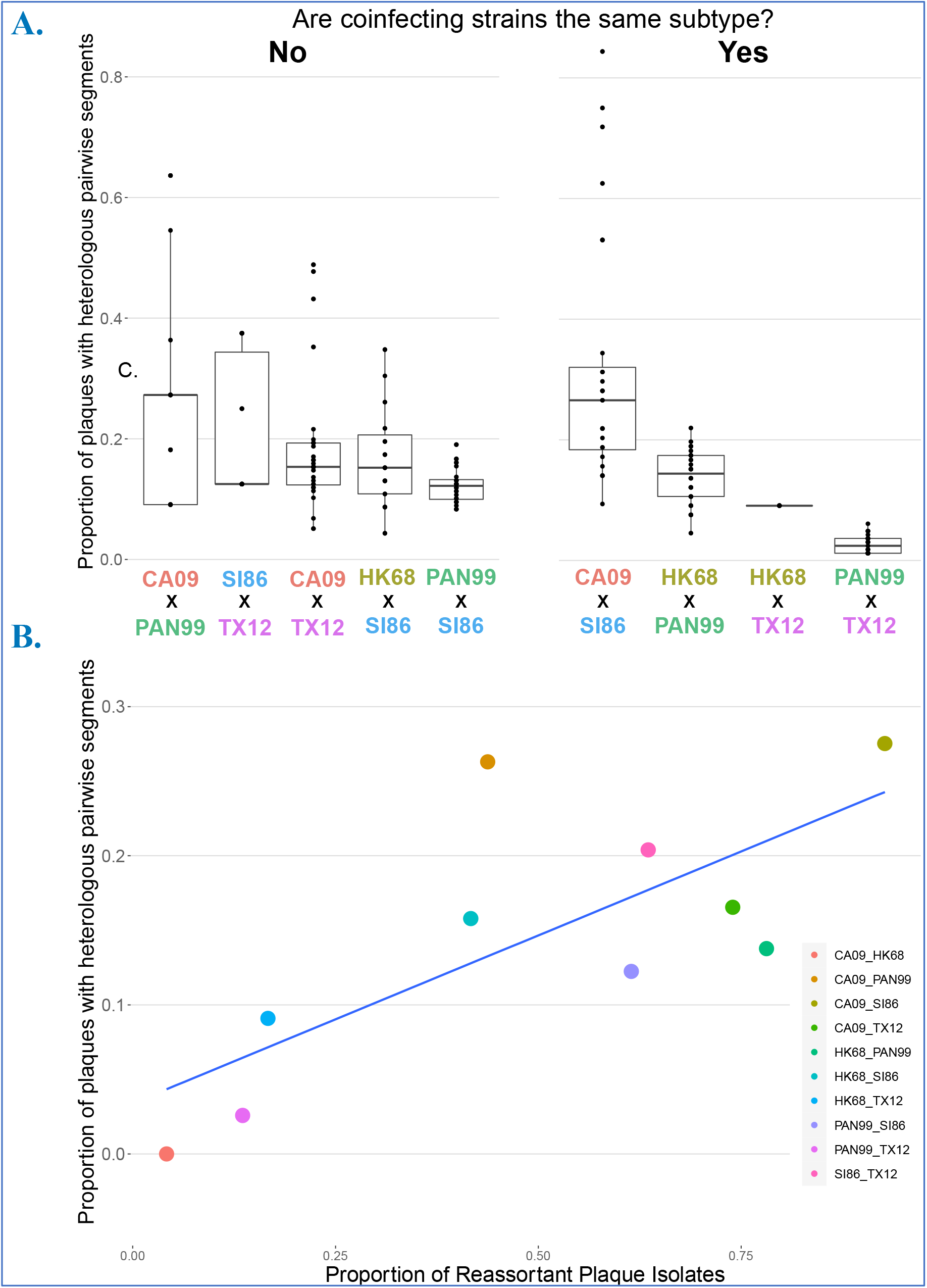
Pairwise segment combinations across experimental coinfections. **A**. Proportion of plaque isolates that had heterologous (i.e. reassortant) genotypes for each pairwise segment combination, represented by points. The left panel shows intersubtype experimental coinfections (i.e. H1N1xH3N2) and the right panel shows intrasubtype (i.e. H1N1xH1N1 or H3N2xH3N2). **B**. Relationship between the reassortment rate (x-axis) and the average proportion of plaque isolates with heterologous genotypes (across all pairwise segment combinations for each experimental coinfection). Each point represents an experimental coinfection.

Homologous combinations in each locus pair composed most progeny genotypes. Overall, across all coinfections, pairwise loci combinations were represented by 53.68% homologous plaques and 15.96% heterologous plaques (see also Figure 7A). This dominance of parental segment associations led to the question of whether this pattern was driven by singly infected cells –which should be very rare under MOI 10, equal proportion infection conditions– or double infections of only parentals. To examine only plaques known to be derived from cells coinfected by different strains, we examined only plaques with reassortant genotypes, thereby discarding progeny produced from potentially singly-infected cells. Considering only reassortant plaques, we still found that homologous pairwise loci combinations dominated among plaques: 42.28% homologous plaques versus 27.21% heterologous plaques. Differences among the strain combinations in experimental coinfections remained (ANOVA: F = 24.269, p < 0.0001) and did not relate to genetic similarity or subtype.

### High reassortment coinfections produce a greater number of unique genotypes

Finally, to examine complete genotypes produced by each experimental coinfection, we determined the eight-segment composition of each plaque and calculated genotype richness (S), the number of unique eight-segment combinations (**Figure 8A**).

**Figure 8.**
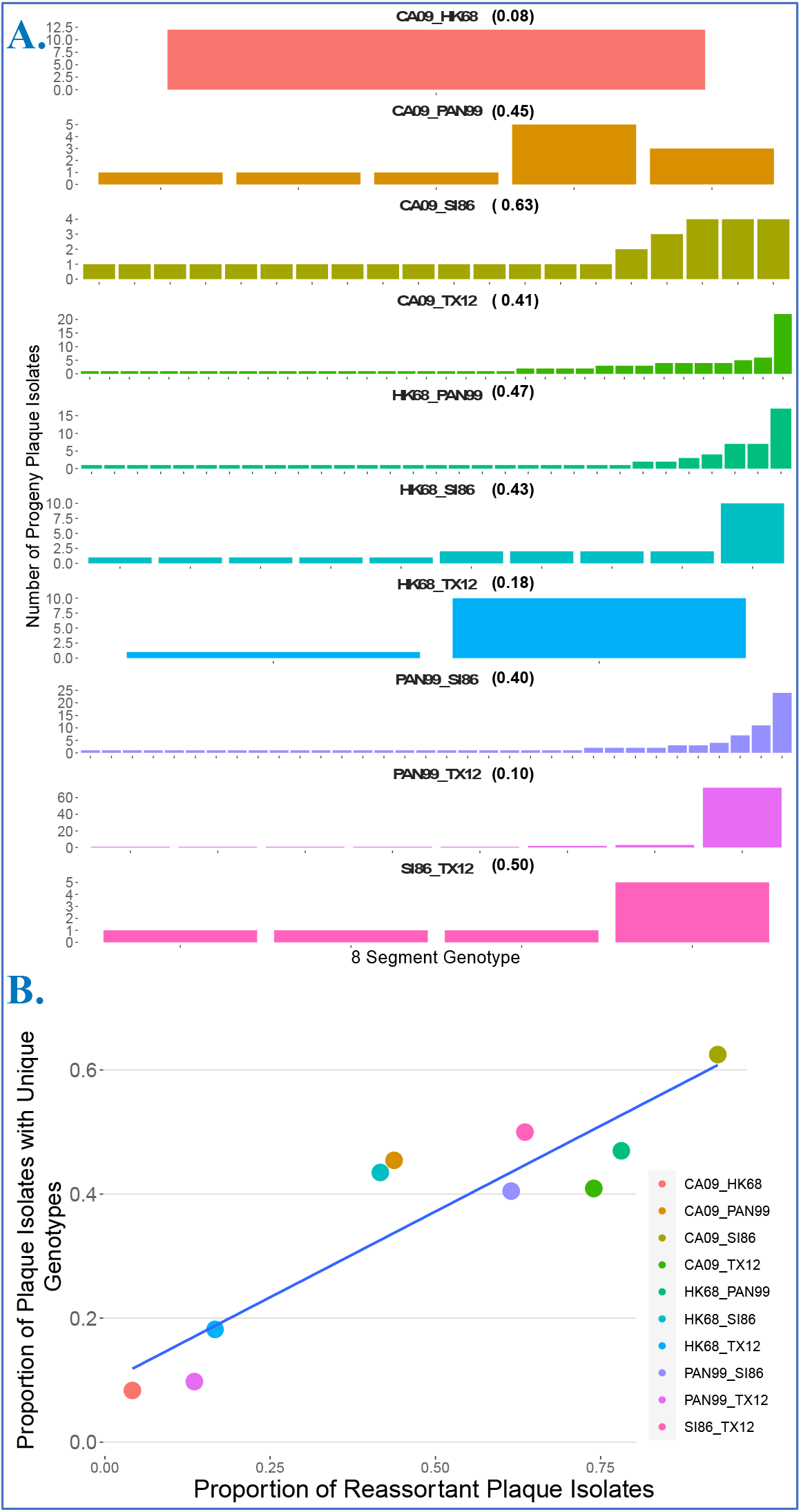
Experimental coinfections generate varying numbers of unique genotypes. **A**. The number of plaques having different 8 segment genotypes are depicted for each experimental coinfection. **B**. The reassortment rate is positively correlated to the proportion of unique genotypes.

Experimental coinfections yielded a maximum of 36 unique genotypes (range: 1-36) from 12-88 isolated plaques with complete eight-segment genotypes (progeny with missing segment data were excluded). The rate of reassortment showed a positive correlation with genotype richness (**Figure 8B**, Multiple R^2^ = 0.8366, p < 0.001).

Because only 94 potential plaques were sampled among 2^8^ = 256 possible combinations (and only a subset of those yielded complete genotypes) we explored the effect of sampling (i.e. how many progeny are genotyped) using a simulation which assumed perfectly random assortment (Bernoulli trial, i.e each progeny isolate has a 0.5 probability of receiving either parental genotype, independently for each segment). One thousand trials of this simulation suggested that sampling 96 plaques would at best yield 90 unique genotypes (mean ± SD = 80.30 ± 3.05). The coinfection that proportionally generated the most unique genotypes, CA09xSI86 (**Figure 9A**, *red point*), yielded 32 plaques (y-axis) with complete genotypes, of which 20 were unique (x-axis). This yield falls outside of the range of 27-32 unique genotypes predicted (blue lines) in a simulation of free assortment for a sample size of 32 plaques. Thus, increased sampling of progeny is necessary to capture rare genotypes: under free reassortment, even 96 sampled progeny would capture ~31.21% of possible 8-segment genotypes (**Figure 9A**)

**Figure 9.**
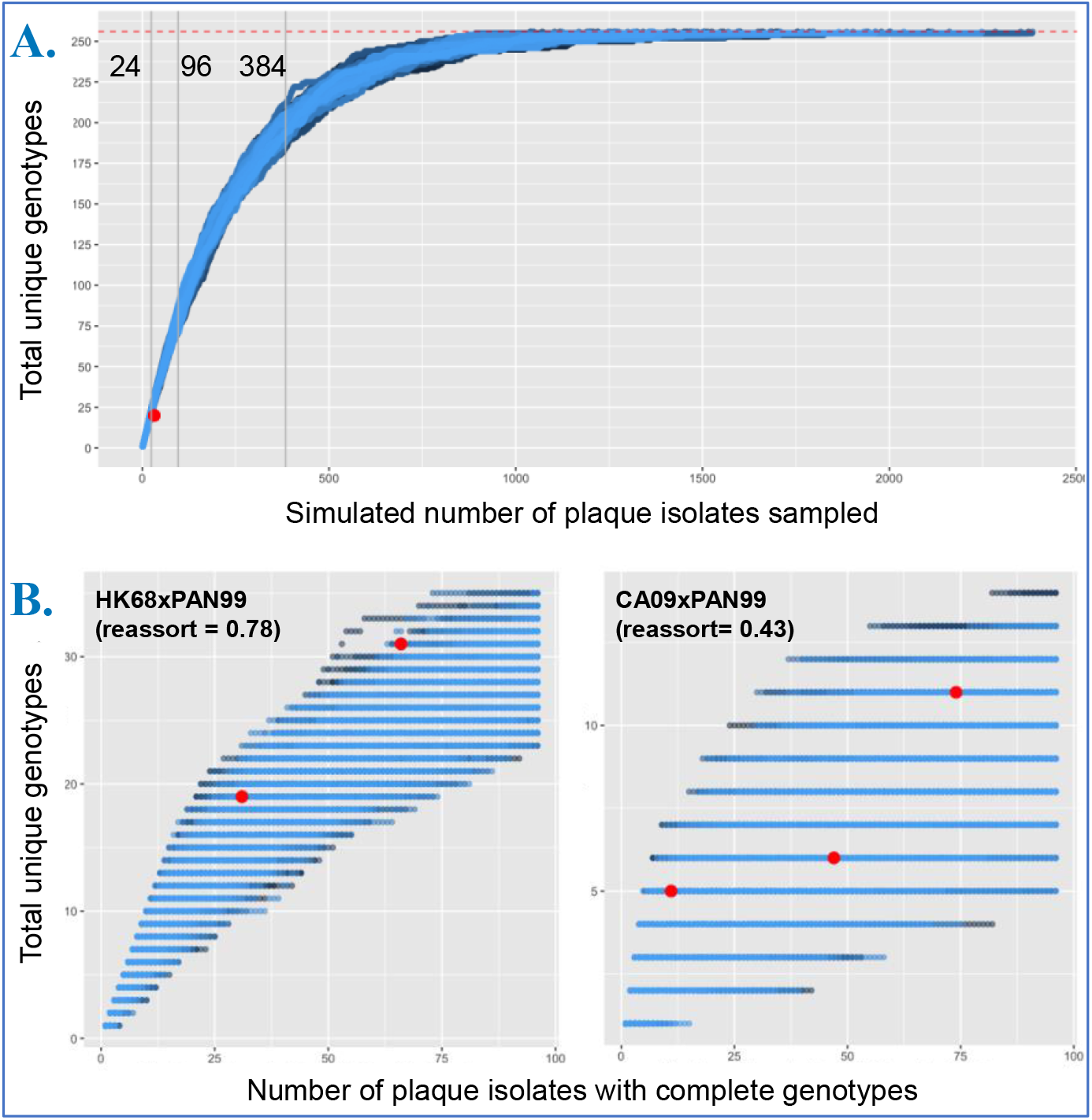
A large sample of progeny plaque isolates is essential to capture all potential genotypes generated by coinfections. **A**. Simulated number of progeny plaque isolates (x-axis) needed to reach 256 genotypes under free reassortment (horizontal red dashed line). Vertical gray lines denote typical sample sizes for molecular biology studies (24, 96, 384). **B**. Replicate coinfection (n = 2, left; n = 3 right) experiments were used as starting data to generate a sample accumulation curve similar to A, but sampling empirical data. Number of unique genotypes found in individual replicate experiments are shown in red, to depict the effect of reduced sample sizes in capturing genotype richness of experimental coinfections.

To achieve sampling all 256 genotypes under random assortment (**Figure 9A**, *dashed red line*), an average of 1,557 ± 333 progeny would have to be sampled and genotyped (n =100 trials, blue lines in **Figure 9A**). However, this result is only expected if the coinfecting strains have free assortment between all segments, which is not the case (Figures 6, 7).

Therefore, to inform on the effect of sampling on estimates of genotype richness and construct a similar genotype accumulation curve, we used biological replicates on a subset of strains. For two replicate HK68xPAN99 coinfections (**Figure 9B**), we screened 188 plaques and obtained a total of 96 complete eight-segment genotypes (x-axis) that yielded 40 unique genotypes (y-axis). If we assume this baseline is the entire genotype space, we can calculate how close each replicate coinfection individually came to capturing all genotypes. In two replicate coinfections, we obtained 77.50% (n = 66 complete genotypes) and 47.50% (n = 30 complete genotypes) of assumed possible eight-segment combinations (**Figure 9B**, *red points*). Similarly, for three replicate CA09xPAN99 coinfections (**Figure 9C**), we screened 282 plaques and obtained a total of 132 complete eight-segment genotypes that yielded only 14 unique genotypes. In our three replicate coinfections we obtained 35.71% (n = 11), 42.86% (n = 47), 78.57% (n = 74) of assumed possible eight-segment combinations (**Figure 9C**, *red points*). These results suggest that obtaining >60 complete genotypes will recover the majority of genotypes in strain combinations that do not reassort freely. For strain combinations that reassort close to freely, robust sample sizes are required to capture all genotypes because of significant sampling variability due purely to chance.

## Discussion

To quantify the outcomes of coinfection between intact strains and their reassortment potential, we employed a high throughput protocol that can examine diverse human seasonal strains spanning decades. Using this robust data set, we were able to draw conclusions regarding strain traits that may influence reassortment potential, and to examine associations between coinfection outcomes that affect reassortment potential. First, we found that the reassortment rate (proportion of reassortants) is an emergent property of a pair of strains and is not explained by strain similarity or shared antigenic subtype. However, we show evidence that strain identity could serve as a predictor as particular strains tend to favor more or less reassortment, independently of the coinfecting partner. Second, we found that most strain combinations involved all 8 segments in reassortment and that random assortment of individual segments among progeny was rare but was not precluded by intersubtype coinfection. Third, we found evidence of non-random pairwise segment associations across most strain combinations, with homologous linkages preferred, but this outcome again was not related to strain similarity or shared subtype. Fourth, we found a range in the number of unique genotypes generated by each coinfection, again suggesting emergent outcomes from distinct strain combinations. Examining associations between coinfection outcomes, we observed that the reassortment rate is closely correlated to both the proportion of unique genotypes and the proportion of progeny with heterologous pairwise segment combinations, but not to segment assortment. Finally, we show that viral kinetics vary independently of the reassortment rate, affecting the total number of reassortants produced by each coinfection.

The highest reported rate of reassortment between human influenza A strains was an average of 88.4% between nearly identical PAN99 strains (Marshall et al. 2013). In comparison our data yielded a 92.71% rate for a CA09xSI86 coinfection (despite the modest sample size, see Figure 9), showing that some divergent strains can potentially generate higher reassortment rates than near-identical strains, albeit, still short of theoretical free reassortment 99.22% = 254/256. This result suggests that strain similarity is not a prerequisite for a high reassortment rate between human influenza strains, as first reported by Phipps et al (Phipps et al., 2017) for heterosubtypic coinfection of H3N2 (PAN99) and H1N1 (A/Netherlands/602/2009(H1N1)-like) strains. Our data further support a reversal of the consensus arising from initial experimental coinfection and vaccine production studies, that coinfection of dissimilar strains or subtype mismatched strains limit the production of reassortant progeny (Essere et al., 2013; Fulvini et al., 2011; Greenbaum et al., 2012; Li et al., 2008).

Thus, overall genetic similarity and subtype do not appear to be consistent predictors of reassortment rate. By testing diverse human seasonal strains, we were able to determine that the reassortment rate appears to be an emergent property of strain combinations. The question was which strain properties are predictive of the reassortment rate. We present initial evidence that strain identity can serve as a predictor of reassortment rate, suggesting that genetic exchange may be an individual trait of influenza A strains, analogous to variation among individuals in meiotic recombination between animals (Coop & Przeworski, 2007, Ritz et al. 2017, Smukowski & Noor 2011). Although genetic exchange clearly requires at least two organisms, these may have mechanisms to predispose to more or less genetic exchange, as characterized in mammals (humans, cattle, and sheep) and *Drosophila* (Ritz et al., 2017). The potential mechanisms that could mediate this trait are plentiful and beyond the scope of this study. Future studies are needed to examine which genotypes and molecular mechanisms underlie the tendency of strains that tend to increase or decrease reassortment.

We found no difference in reassortment rate between intrasubtype and intersubtype strain pairings. Influenza A H3N2 and H1N1 subtypes have co-circulated in the human population for nearly half a century, with H1N1 1918 pandemic lineage strains since 1977 (Zhdanov et al., 1978) and 2009 pandemic lineage strains (H1N1pdm) since that year. Given their co-circulation, it may be expected that H1N2 reassortants would be detected. Indeed, H1N2 reassortants have been identified (Chen et al., 2006; Mukherjee et al., 2012; Nelson et al., 2008b) and have given rise to epidemics (Gregory et al., 2002) or become epidemiologically significant (Nelson et al., 2008a). However, H1N2 strains have not managed to establish themselves in the human population (Chen et al., 2006; Gregory et al., 2002). Our results imply that the barrier for *generating* H1N2 reassortants might actually be low and thus —given that some H1N2 strains have generated epidemics— the reason why they have not become established likely relates to competition between subtypes in circulation, and not to constraints on reassortment. One limitation of our study is that we could not statistically determine if *individual* subtypes differed in reassortment rate (i.e. H1N1/H1N1 versus H3N2/H3N2 coinfections), due to sample size constrains. This may be important data to forecast the risk of immune evasion due to intrasubtypic reassortment in seasons where H1N1 or H3N2 subtypes dominate.

Even in the coinfections with the highest reassortment rates, segment exchange was not free. Overall, there was a bias towards specific parental segments among the progeny homologous pairwise combinations in reassortants. This finding is in line with numerous experimental coinfection studies, dating from the first study of this type (Essere et al., 2013; Greenbaum et al., 2012; Li et al., 2008; Lubeck et al., 1979; Phipps et al., 2017). Our novel finding is that the pattern of non-random segment exchange is not mediated by the genetic distance of strains involved or intersubtypic combinations, as widespread consensus in the literature held (Essere et al., 2013; Li et al., 2008; Lubeck et al., 1979). One of the strains that showed the freest segment assortment among progeny (Figure 6B: 5/8 segments assorting randomly) was SI86xPAN99, an intersubtype coinfection. Likewise, the strain with the most heterologous pairwise segment combinations was SI86xCA09, from divergent H1N1 lineages. These findings within human strains echo other studies that showed robust, if not random, segment exchange among distantly related strains (Li et al., 2010; Octaviani et al., 2010).

Although we did not find strain combinations that approached random assortment between segments, we did find that there was a tight and statistically significant correlation between reassortment rate and both the proportion of unique genotypes and the proportion of heterologous pairwise segment combinations (these two are most likely tightly correlated). While *prima fasciae* this seems like a trivial observation, there is no *a priori* reason why these should be linked; for example, in the extreme case of a single favorable heterologous pairing of two segments, there could be only one genotype produced and thus 100% reassortant progeny. If the above correlation holds, the implication is that strain combinations that yield a higher rate of reassortants also produce more diverse combinations.

Another important relationship between coinfection outcomes we found is that viral kinetics vary independent of the reassortment rate, affecting the total number of reassortants produced by each coinfection. Reassortment studies typically focus on rates and proportions among progeny. However, an increasing number of studies show that coinfection itself can affect influenza A virus growth kinetics (Jacobs et al., 2019; Martin et al., 2020; Phipps et al., 2020; Rimmelzwaan et al., 1998), a long-held observation from vaccine production studies (Frensing et al., 2013; Isken et al., 2012). In single strain infections this phenomenon has been termed multiplicity dependence (Phipps et al., 2020). We therefore reasoned that mixed coinfection (at our high MOI experimental conditions) could impact viral production. We find that highly productive infections do not have a higher rate of reassortment (and vice versa), but productivity will have an impact on the total number of reassortants in all cases. The lack of correlation between viral production and reassortment rate could assist in public health risk assessment by using these traits as separate criteria when assessing reassortment potential.

A last technical observation is that data arising from simulations and empirical data suggest that it is necessary to sample a large number of progeny plaque isolates to cover the potential diversity generated by each coinfection and to avoid random sampling variability (due purely to chance). This observation is purely statistical and this limitation applies to the current study. However, provided complete genotypes are obtained, 96 samples will provide most unique genotypes, at least in strain combinations that do not have completely free segment exchange.

An important study design and conceptual element regarding this work bears discussion, as it has implications for data analysis and interpretation of all studies examining reassortment: we sought to examine the outcomes of coinfection of intact strains and any reassortment potential. We focused on maintaining unbiased conditions that examined a single cycle of replication (Marshall et al., 2013). By design we avoid making inferences regarding the fitness of progeny viruses (beyond capacity to initiate infection in plaque assay). As indicated in the first experimental coinfection study (Lubeck et al., 1979), it is important to analyze reassortants arising from a single cycle of replication, as multicycle growth may enrich for particular isolates, due to growth competition among progeny unrelated to the intracellular process of reassortment. Furthermore, when assaying viral “fitness” in the lab, the conclusions are applicable only to those conditions, which may or may not be representative of conditions in nature. A limitation of our study design in determing reassortment potential is that we cannot distinguish between solo infections (which are expected to be rare at MOI = 10) or double infections of the same strain in a cell. Distinguishing those outcomes at the single cell level is technically challenging, and we will embark on this objective in future studies. In particular, one cannot assume that progeny viruses with parental genotypes arise only from cells infected with one strain; they may simply not reassort. Our goal was to examine the overall outcome of coinfection of intact strains to reveal areas for future investigation.

At an evolutionary level, our data suggest that viral genetic exchange is potentially an individual social trait subject to natural selection (Díaz-Muñoz et al. 2017). A number of papers in influenza (Brooke, 2017; Koelle et al., 2019; Phipps et al., 2020) and other segmented viruses (Díaz-Muñoz et al., 2013; Turner and Chao, 1999, 1998) have suggested this possibility, and here we provide experimental evidence to more firmly establish this conclusion. This conclusion suggests some directions for such future experiments. Segment mismatch studies should include more segments, as protein compatibility may be strain dependent and protein incompatibilities (e.g. among polymerase complex subunits) can be less severe among divergent strains (Octaviani et al. 2010) than more closely related strains (Li et al. 2008, Octaviani et a. 2011). At an extreme, a study of influenza A & B reassortment has even shown successful intertypic expression of the HA segment (Baker et al. 2014). For studies of RNA-RNA interactions and viral assembly, our data support the idea that RNA-RNA interaction networks among segments are not conserved (Gavazzi et al. 2013, Lakdawala et al. 2014), and may retain flexibility for interactions with different strains (Le Sage et al. 2020). Finally, for most studies of coinfection, our data suggests that conclusions based on one or two strains should be extrapolated cautiously. For instance, intersubtype reassortment is often assumed to be less pervasive than intrasubtype reassortment (Nelson et al., 2008a) and our data suggests weak evidence for this assumption (Figure 3), or at a minimum, that there will be some exceptions.

The evolutionary observation of emergent and strain dependent reassortment also has some important public health implications. First, the tendency of particular strains to increase reassortment rate (and potentially generate more unique genotypes) implies that the risk of reassortment is not evenly spread among strains. Identification of these high-reassortment strains could improve pandemic preparedness or reveal new treatment targets, particularly if specific loci can be used as markers for increased genetic exchange. One example that has been discussed in the literature is avian H9N2 viruses which tend to provide backbones for avian H7N9s (Pu et al., 2021, 2015), creating a highly reassortant platform that is responsible for recent human infections (Chen et al., 2013; Gao et al., 2013). Second, exchange of a subset of segments even at a low frequency (e.g. SI86xHK86, which primarily exchanges HA, PA, and NP segments), could be potentially important in the emergence of novel strains under the correct selective regimes, particularly if those segments carry virulence or host tropism determinants. Thus, we believe for pandemic preparedness our base assumption should be that in virtually every coinfection influenza viruses are shuffling the deck, not perfectly but constantly.

## Acknowledgements

This project was funded by NIH grant 4R00AI119401-02 to SDM. SDM was also supported by an NIH Pathway to Independence Fellowship (grant 1K99AI119401-01A1). KYT and IA were supported by NIH grant 4R00AI119401-02 to SDM. Ted Ross kindly provided strains from his collection. SM was supported by the Academy of Finland Postdoctoral Fellowship (grant no. 323426). Graham Coop and Daniel Runcie provided feedback on specific analyses. Anice Lowen, Chris Brooke, and Seema Lakdawala provided feedback on the manuscript.

## Notes

### Competing Interest Statement

The authors have declared no competing interest.

