## Supplementary Materials for "Influenza A virus reassortment is strain dependent"

### SUPPLEMENATRY MATERIALS

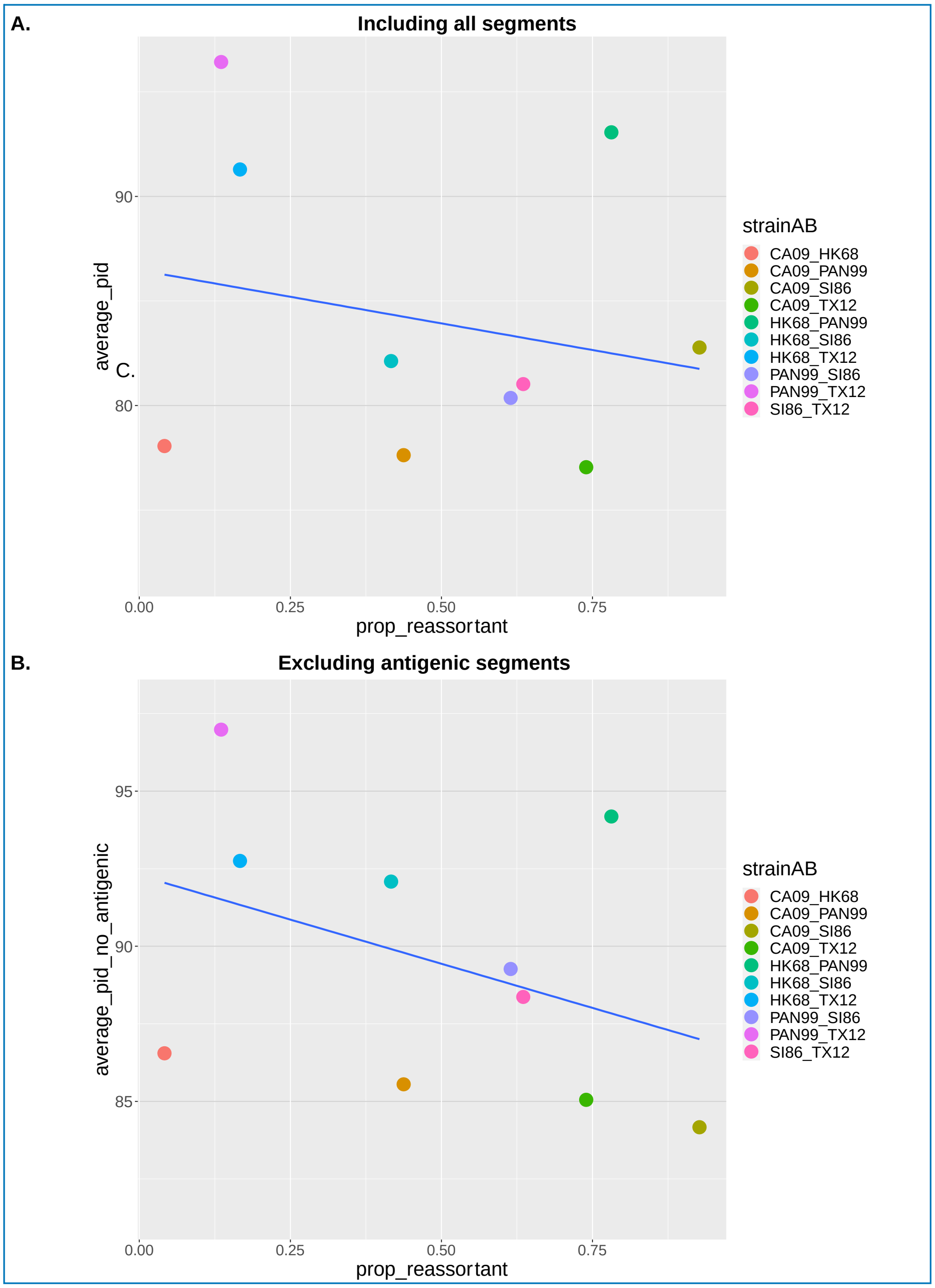


**Figure S1.** Correlation between reassortment rate and genetic similarity between strains in each experimental coinfection. **A.** Genetic similarity calculated using all eight segments. **B.** Genetic similarity calculated excluding antigenic segments HA and NA.

| Strain | **CA09** | **HK68** | **PAN99** | **SI86** | **TX12** | Segment |
|---|---|---|---|---|---|---|
| CA09 | | | | | | PB2 |
| HK68 | 89.1 | | | | | PB2 |
| PAN99 | 84.0 | 94.2 | | | | PB2 |
| SI86 | 85.1 | 91.3 | 89.2 | | | PB2 |
| TX12 | 83.9 | 92.7 | 97.1 | 88.5 | | PB2 |

| Strain | CA09 | HK68 | PAN99 | SI86 | TX12 | |
|---|---|---|---|---|---|---|
| CA09 | | | | | | PB1 |
| HK68 | 91.3 | | | | | PB1 |
| PAN99 | 94.7 | 94.1 | | | | PB1 |
| SI86 | 81.4 | 82.7 | 81.7 | | | PB1 |
| TX12 | 93.0 | 92.4 | 96.7 | 81.1 | | PB1 |

| Strain | CA09 | HK68 | PAN99 | SI86 | TX12 | |
|---|---|---|---|---|---|---|
| CA09 | | | | | | PA |
| HK68 | 82.9 | | | | | PA |
| PAN99 | 83.3 | 93.4 | | | | PA |
| SI86 | 83.1 | 94.4 | 91.3 | | | PA |
| TX12 | 83.3 | 92.2 | 97.0 | 90.3 | | PA |

| Strain | CA09 | HK68 | PAN99 | SI86 | TX12 | |
|---|---|---|---|---|---|---|
| CA09 | | | | | | HA |
| HK68 | 53.2 | | | | | HA |
| PAN99 | 54.4 | 89.2 | | | | HA |
| SI86 | 77.3 | 53.8 | 54.9 | | | HA |
| TX12 | 53.9 | 85.9 | 94.4 | 66.9 | | HA |

| Strain | CA09 | HK68 | PAN99 | SI86 | TX12 | |
|---|---|---|---|---|---|---|
| CA09 | | | | | | NP |
| HK68 | 84.7 | | | | | NP |
| PAN99 | 82.9 | 92.9 | | | | NP |
| SI86 | 84.6 | 94.2 | 90.3 | | | NP |
| TX12 | 82.2 | 91.6 | 96.7 | 89.1 | | NP |

| Strain | CA09 | HK68 | PAN99 | SI86 | TX12 | |
|---|---|---|---|---|---|---|
| CA09 | | | | | | NA |
| HK68 | 52.0 | | | | | NA |
| PAN99 | 53.3 | 90.2 | | | | NA |
| SI86 | 79.9 | 50.7 | 52.4 | | | NA |
| TX12 | 52.2 | 87.9 | 95.1 | 51.1 | | NA |

| Strain | CA09 | HK68 | PAN99 | SI86 | TX12 | |
|---|---|---|---|---|---|---|
| CA09 | | | | | | M |
| HK68 | 88.8 | | | | | M |
| PAN99 | 86.9 | 96.3 | | | | M |
| SI86 | 88.1 | 95.7 | 93.6 | | | M |
| TX12 | 86.7 | 94.6 | 97.3 | 92.4 | | M |

| Strain | CA09 | HK68 | PAN99 | SI86 | TX12 | |
|---|---|---|---|---|---|---|
| CA09 | | | | | | NS |
| HK68 | 82.5 | | | | | NS |
| PAN99 | 81.5 | 94.2 | | | | NS |
| SI86 | 82.7 | 94.2 | 89.5 | | | NS |
| TX12 | 81.2 | 93.0 | 97.1 | 88.8 | | NS |

**Table S1.** Pairwise nucleotide identity between strains used in experimental coinfections by segment.

| **Strains in Coinfection** | **Total Segments** | **Mean proportion** | **Std Dev proportion** | **Number of Segments that assort randomly** |
|---|---|---|---|---|
| **HK68_TX12** | 8 | 0.967 | 0.0289 | 0 |
| **CA09_PAN99** | 8 | 0.888 | 0.147 | 1 |
| **CA09_TX12** | 8 | 0.708 | 0.138 | 2 |
| **PAN99_SI86** | 8 | 0.615 | 0.0839 | 6 |
| **CA09_SI86** | 8 | 0.727 | 0.118 | 1 |
| **SI86_TX12** | 8 | 0.825 | 0.131 | 1 |
| **CA09_HK68** | 8 | 0.993 | 0.0115 | 0 |
| **HK68_PAN99** | 8 | 0.757 | 0.131 | 2 |
| **PAN99_TX12** | 8 | 0.968 | 0.0221 | 0 |
| **HK68_SI86** | 8 | 0.895 | 0.0778 | 0 |

**Table S2.** Representation of segments among progeny plaque isolates for each experimental coinfection. Each genotype entered the coinfection at a 0.5 proportion. For each coinfection, across all segments, the mean proportion of each strain genotype is shown. The number of segments that show random assortment (i.e. fall within the confidence interval for a 50:50 ratio) are listed in the last column.

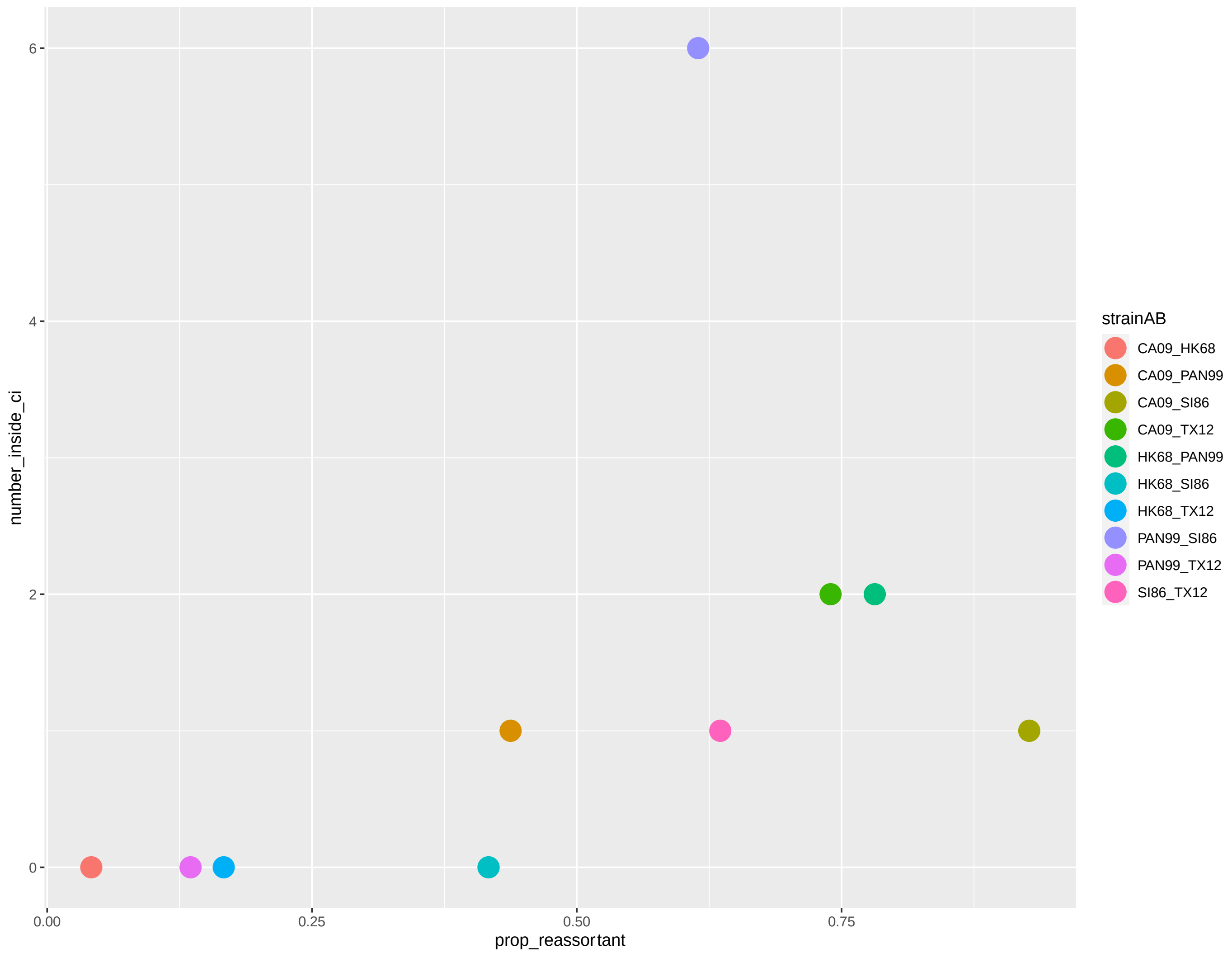


**Figure S2.** Relationship between the reassortment rate and the number of segments that individually assort randomly among the progeny.

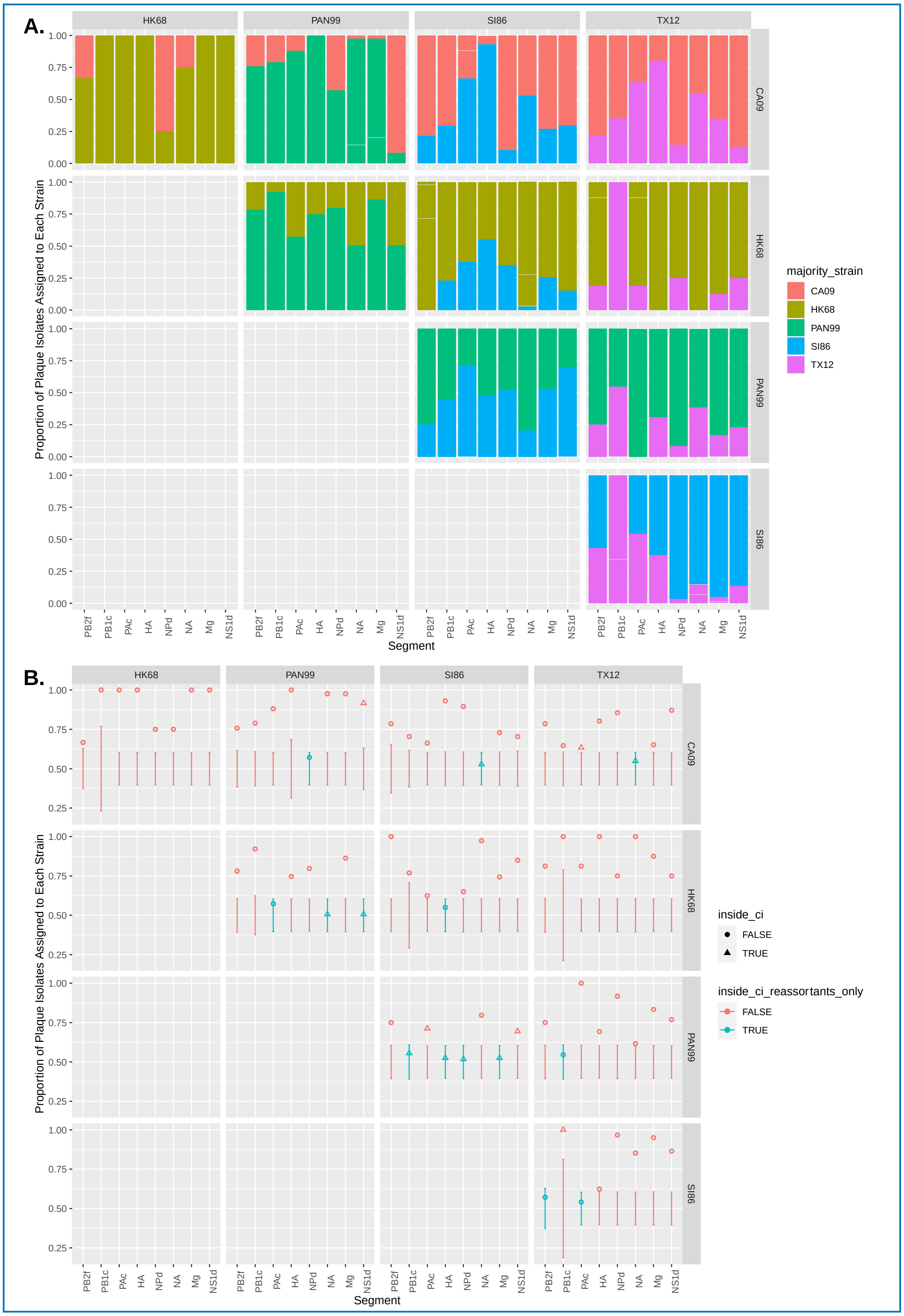


**Figure S3.** Segment representation in progeny plaque isolates with regard to strain, using only reassortant plaques. **A.** Plot shows frequency of each strain's allele for each segment in plaque isolates. **B.** Depicts which segment frequencies are within (blue points) or outside (red points) the confidence interval for 50:50 distribution of strain alleles among plaque isolates.

#### Detailed Genotype by Barcode Sequencing (GbBSeq) Methods

*Amplicon design and analysis strategy*

We designed amplicons for each of the eight segments: for antigenic proteins, which show high sequence variability we designed two primers, one for each human influenza virus subtype for a total of 10 loci.

The amplicons were generated by priming to the uni13 at the 5' end of influenza vRNAs, which is shared among all influenza A strains (Zhou et al., 2009) and to an internal region specific to each segment.

*Barcode design*

Barcodes were selected using the approach of Buschmann and Bystrykh (Buschmann et al., 2014; Buschmann and Bystrykh, 2013) as implemented in the R package DNABarcodes. Eight base pair barcodes were generated to maximize the Levenshtein distances (d = 3, Sequence-Levenshtein in paper), to guarantee correction of at least one error (substitution, insertion, deletion). Because the Illumina sequencing platform does not have trouble with triplet homopolymers, these were allowed in the barcodes. However, barcodes containing the GGC motif, which is known to be associated with sequencing errors in this platform, were not selected.
